# Mouse Genome-Wide Association and Systems Genetics Identifies *Lhfp* as a Regulator of Bone Mass

**DOI:** 10.1101/454678

**Authors:** Larry D. Mesner, Gina Calabrese, Basel Al-Barghouthi, Daniel M. Gatti, John P. Sundberg, Gary A. Churchill, Dana. A. Godfrey, Cheryl L. Ackert-Bicknell, Charles R. Farber

## Abstract

Bone mineral density (BMD) is a strong predictor of osteoporotic fracture. It is also one of the most heritable disease-associated quantitative traits. As a result, there has been considerable effort focused on dissecting its genetic basis. Here, we performed a genome-wide association study (GWAS) in a panel of inbred strains to identify associations influencing BMD. This analysis identified a significant (P=3.1 x 10^−12^) BMD locus on Chromosome 3@52.5 Mbp that replicated in two seperate inbred strain panels and overlapped a BMD quantitative trait locus (QTL) previously identified in a F2 intercross. The association mapped to a 300 Kbp region containing four genes; *Gm2447, Gm20750, Cog6*, and *Lhfp*. Further analysis found that Lipoma HMGIC Fusion Partner (*Lhfp*) was highly expressed in bone and osteoblasts and its expression was regulated by local expression QTL (eQTL) in multiple tissues. A co-expression network analysis revealed that *Lhfp* was strongly connected to genes involved in osteoblast differentiation. To directly evaluate its role in bone, *Lhfp* deficient mice (*Lhfp*^−/−^) were created using CRISPR/Cas9. Consistent with genetic and network predictions, bone marrow stromal cells (BMSCs) from *Lhfp*^−/−^ displayed increased osteogenic differentiation. *Lfhp*^−/−^ mice also had elevated BMD due to increased cortical bone mass. In conclusion, we used GWAS and systems genetics in mice to identify *Lhfp* as a regulator of osteoblast activity and bone mass.

## INTRODUCTION

It is currently estimated that half of all Americans over the age of 50 already have or are at high risk of developing osteoporosis [1]. Bone mineral density (BMD) is used clinically to diagnose osteoporosis and beyond age, it is the single strongest predictor of the risk of fracture [2]. BMD is also one of the most heritable disease-associated quantitative traits with studies demonstrating that up to 80% of the variance in peak bone mass is heritable [3–6]. Consistent with its high heritability, genome wide association studies (GWASs) in humans have identified hundreds of loci for BMD [7–9]. However, only a small fraction of the variance in BMD can be collectively explained by these loci, suggesting that BMD is influenced by a large number of small effect size loci [10]. As a result, there remains much to be discovered regarding the genetics of bone mass and genetic mapping efforts using mouse models is a complementary approach to identify novel regulators of bone mass [11–13].

Historically, linkage analyses in intercrosses, backcrosses and recombinant inbred strain panels were the mainstay of mouse genetics [14]. These approaches were used to identify dozens of quantitative trait loci (QTL) for BMD and other bone traits [15,16]. However, identifying causative genes underlying QTL proved challenging [17]. Over the last decade, gene mapping approaches have transitioned from low-resolution linkage mapping to high-resolution GWASs [11]. The first GWASs in mice used panels of inbred mouse strains [18–21] and by leveraging the accumulation of recombinations across generations, this approach significantly increased mapping resolution [19]. However, the approach was limited by population structure and low statistical power, due to the complicated breeding histories of inbred mouse strains and small number of easily accessible and appropriate inbred strains (N typically < 30), respectively. Later studies demonstrated that these issues could be partly addressed by accounting for population structure and leveraging information from linkage-based QTL studies [22–24]. Given the significant amount of existing phenotypic and genotypic data on inbred strain panels [25], this approach is potentially a cost-effective strategy to identify novel regulators of complex traits.

High-resolution mapping approaches have significantly increased our ability to identify narrow regions of the genome harboring trait associated genetic variants. It is still, however, a challenge to go from association to gene to function and several approaches have been developed that can assist in bridging this gap. Specifically, systems genetics approaches involving the integration of other types of “-omics” data have proven useful [26]. Two systems genetics approaches for informing GWAS are expression quantitative trait loci (eQTL) discovery and co-expression network analysis [27]. EQTL discovery allows one to link variants associatied with a trait, such as BMD, to changes in gene expression which leads to the hypothesis that the change in gene expression is causal for the change in phenotype. EQTL studies have been tremendously successful in identifying target genes downstream of genome-wide significant variants (as examples; [28,29]). In many cases, the identified target genes have no known connection to the phenotype under investigation. It has been shown that co-expressed genes often operate in the same pathway or are functionally related [30]. Therefore, by using co-expression networks, which cluster genes based on patterns of co-expression across a series of perturbations [31], it is possible to develop hypotheses as to the function of a novel gene. When a locus has been resolved down to a small number of genes using genetics methods, unknown or poorly characterized genes can be ranked as the most likely candidate based on their function predicted from a co-expression network generated in a disease relevant tissue or cell-type [12,32].

Here, we used GWAS in an inbred strain panel to identify two chromosomal regions harboring variants influencing BMD. One of the associations, located on Chromosome (Chr.) 3, affected BMD in both sexes and was replicated in two seperate inbred strain panels and an F2 intercross. This locus mapped to a 300 Kbp interval (NCBI37/mm9; Chr3:52.5-52.8 Mbp) encompassing four genes, *Gm2447, Gm20750, Cog6*, and *Lhfp*. An eQTL analysis and examination of a bone co-expression network suggested that *Lhfp* was a causal gene at this locus. The analysis of BMD, and other bone parameters, in *Lhfp* mutant mice supported this hypothesis. Thus, we have used GWAS and systems genetics in the mouse to identify *Lhfp* as a novel regulator of bone mass.

## RESULTS

### Identification of genome-wide associations for BMD

We performed a GWAS for total body BMD in 27 classical (non wild-derived) inbred strains at 12 months of age fed a chow diet. Genome scans were performed separately for each sex (**Supplemental Files 1 and 2**). In female mice, a significant (−log10(P)>6) association was identified on Chromosome (Chr.) 3 and, in males, significant (−log10(P)>5.9) loci were identified on Chrs. 2 and 3 (Figure 1).

**Figure 1.**
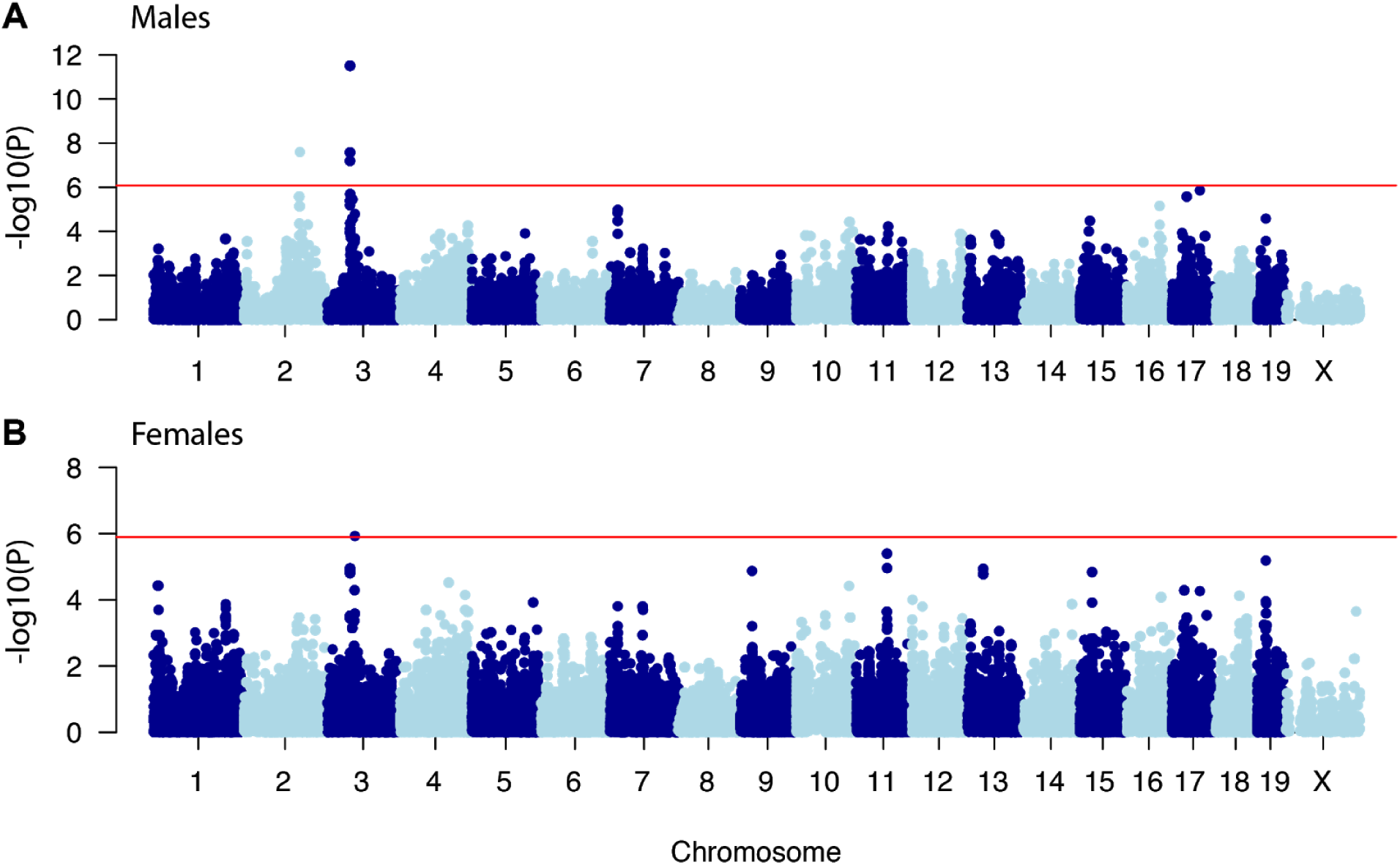
**Manhattan plot for BMD GWAS in the “Ackert” inbred strain panel.** A) GWAS results in male mice. B) GWAS results in female mice. Data from 27 classical inbred strains was used in the GWAS.

### Replication of the Chr. 3 association

Given our goal of identifying novel genes influencing BMD, we selected the Chr. 3 locus for further investigation. This locus was chosen because it was the most significant and the only one identified in both sexes (Figure 1). However, upon closer inspection, Chr. 3 harbored two associations, with peaks at 52.5 and 63.3 Mbp. In males, the 52.5 Mbp peak was the most significant (−log10(P)=11.5), whereas in females the 63.3 Mbp peak was the most significant (−log10(P)=5.9). The lead SNPs at both peaks were in moderate linkage disequilibrium (r^2^=0.46), making it unclear if they represented independent loci. We performed conditional analyses in males and in both cases each peak still exceeded chromosome-wise significance (−log10(P)>2.9) after controlling for the other, suggesting they represent independent loci.

We next sought to identify independent datasets supporting the validity of the Chr. 3 associations. We previously identified a QTL, *Bmd40*, affecting femoral BMD on Chr. 3 in 32 week-old mice from a C57BL/6J (B6) x C3H/HeJ (C3H) (BXH) F2 intercross fed a high-fat diet [16]. The peak of *Bmd40* overlaps both associations (Figure 2A). Importantly, the lead SNPs of the Chr.3@52.5 Mbp association are polymorphic between B6 and C3H. We also identified two sets of inbred strains with total body BMD measurements (the “Naggert” and “Tordoff” studies; data available from the Mouse Phenome Database [25] (https://phenome.jax.org/)) that were large enough (N strains > 25) to attempt to replicate the associations. In both “Naggert” and “Tordoff” panels, the strains used largely overlapped, but they did represent independent measures of BMD at different ages and conditions (Naggert - 15-17 wks old, high-fat diet; Tordoff - 14-18 weeks, chow diet). In both strain sets the exact same sets of SNPs at 52.5 Mbp reached chromosome-wide significance (−log10(P)>2.9) in both sexes (Figures 2B-E). The association at 63.3 Mbp did not replicate in either strain set (Figures 2B-E). These data provide additional support for the BMD association at 52.5 Mbp. Importantly, in all three inbred strain panels (“Ackert”, “Naggert” and “Tordoff”) and the BXH F2 intercross, reference (B6) alleles were associated with increased BMD relative to non-reference (C3H) alleles (Figure 2F). Together, these data, from independent sources, are consistent with the hypothesis that a variant(s) in proximity of 52.5 Mbp on Chr. 3 influences BMD.

**Figure 2.**
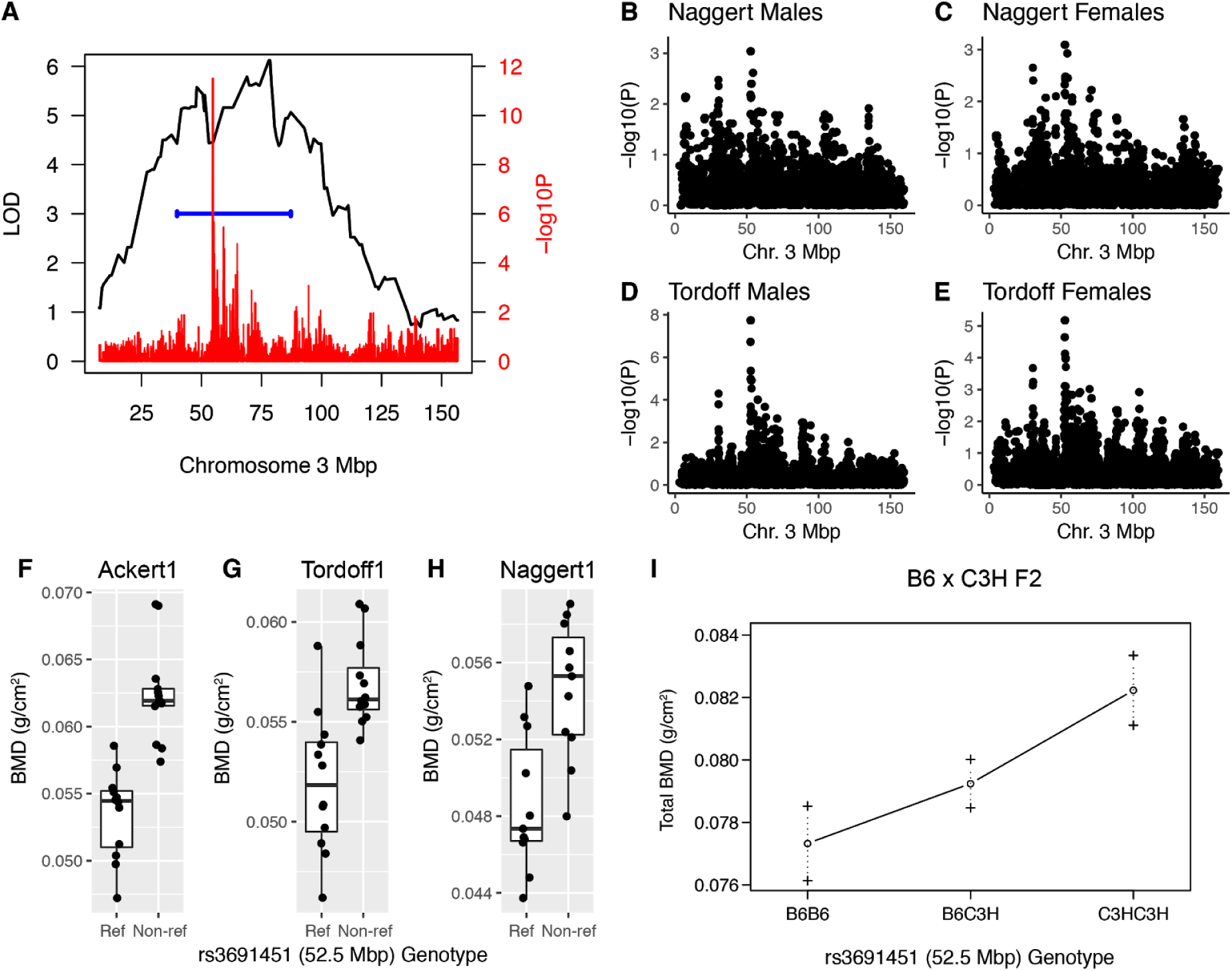
**Replication of BMD association on Chr. 3@52.5 Mbp in both sexes in multiple independent populations.** A) *Bmd40*, a QTL impacting BMD in an F2 cross between C57BL/6J and C3H/HeJ, overlaps the Chr. 3 association at 52.5 Mbp. B-D) Replication of the Chr. 3 association at 52.5 Mbp in two (“Tordoff” (N=30) and “Naggert” (N=31)) different sets of inbred strains. Chromosome-wide significance was −logP>2.9 in both strain sets. F-I) Non-ref (C3H/HeJ) alleles of associated SNPs increase BMD in all four populations.

### The Chr. 3 association implicates a 300 Kbp interval encompassing four transcripts

The set of SNPs that were the most significantly associated with BMD spanned a 300 Kbp interval from 52.5 to 52.8 Mbp (Figure 3A). This region contained four RefSeq transcripts: *Gm2447, Gm20750, Cog6*, and the 5’ end of *Lhfp*. *Gm2447* and *Gm20750* were listed as “predicted” RefSeq transcripts and annotated as long non-coding RNAs (lncRNAs). The evidence for these transcripts was based on prediction models and a small number of expressed sequence tag (EST) sequences. Neither of these transcripts have homologs in humans, rats, or any other mammalian species. To determine if *Gm2447* and *Gm20750* were expressed in mouse bone or bone cells, we performed total RNA-seq (poly A+ and poly A-) on three bone and three marrow-derived osteoblasts samples. *Gm2447* and *Gm20750* were not expressed, whereas the other two transcripts, *Cog6* and *Lhfp*, which are well-annotated protein-coding sequences, were highly expressed in both bone and osteoblasts (Figure 3B). We also analyzed the expression profiles of *Cog6* and *Lhfp* in 96 mouse tissues and cell lines using data available from BioGPS (http://biogps.org/) [33]. *Cog6* was highly expressed in all tissues profiled (Figure 3C). *Lhfp* showed a more restrictive expression profile (Figure 3C). Importantly, *Lhfp* expression in primary calvarial osteoblasts was among the highest of any of the 96 samples surveyed (Figure 3C). *Cog6* is part of the conserved oligomeric Golgi complex required for maintaining normal structure and activity of the Golgi apparatus [34]. *Lhfp* is a member of the lipoma HMGIC fusion partner (*LHFP*) gene family with no known function [35]. All other transcripts on either side of the region were >200 Kbp away.

**Figure 3.**
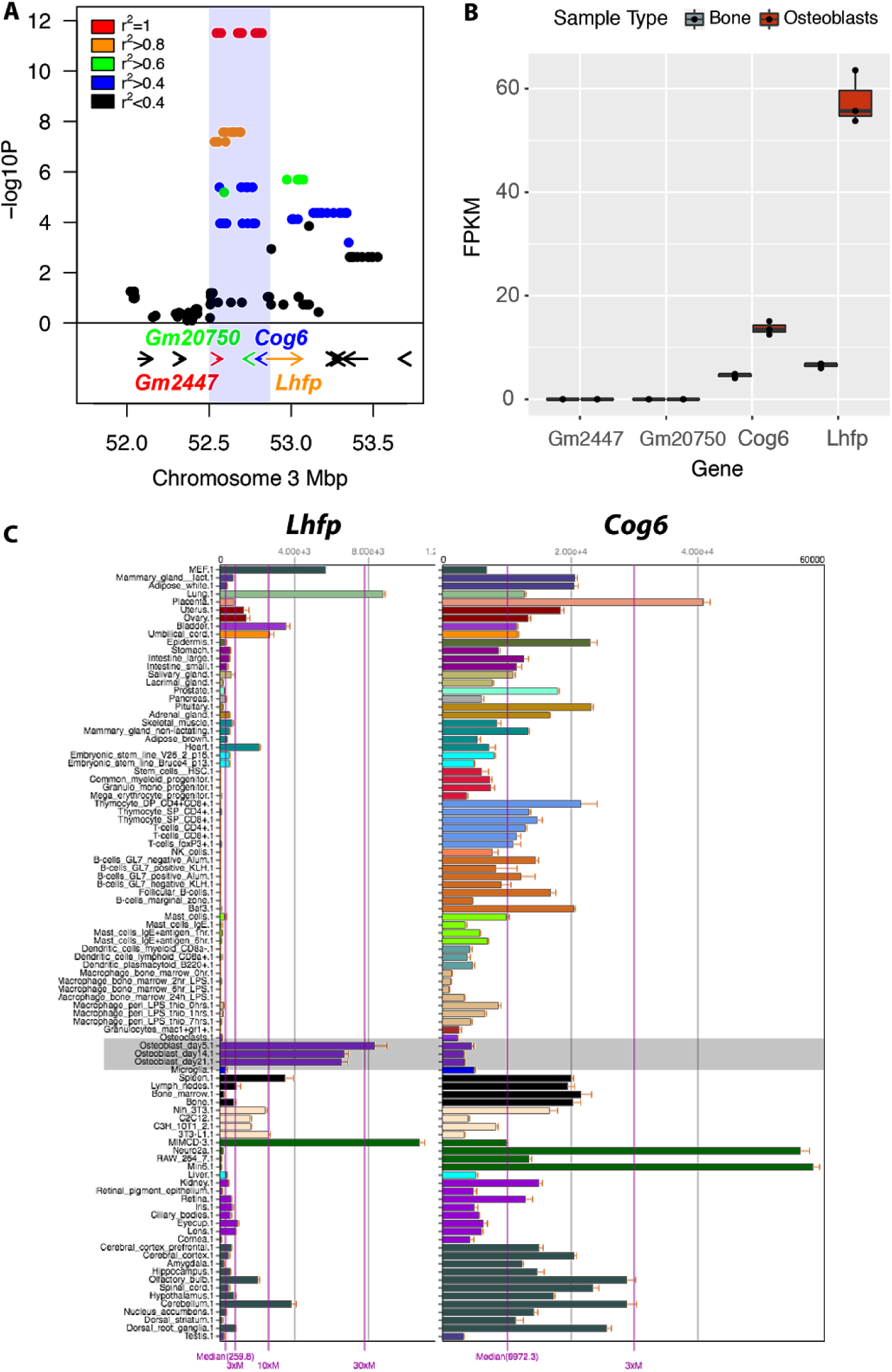
**Interrogation of the Chr. 3@52.5 Mbp association.** A) The association implicates four genes (*Gm2447, Gm20750, Cog6* and *Lhfp*) based on their proximity in the locus. B) RNA-seq expression profiles of the four genes in mouse bone (minus marrow) and osteoblasts derived from bone marrow stromal cells (N=3 each sample type). C) Microarray expression profiles for *Cog6* and *Lhfp* in 96 diverse mouse tissues and cell-types (data from BioGPS, http://biogps.org/) [33].

### Coding polymorphisms in Cog6 and Lhfp

We cannot exclude *Gm2447* and *Gm20750* (or for that matter other genes flanking the association); however, based on the data above we focused on interrogating *Cog6* or *Lhfp* as potential causal genes. First, we evaluated *Cog6* and *Lhfp* for coding polymorphisms among inbred strains. Based on whole genome-sequence data from C57BL6/J and C3H/HeJ (which carry alternative alleles at the association) there are no coding variants between the strains for *Lhfp [36]*. In contrast, there were three non-synonymous SNPs in *Cog6* between B6 and C3H. These SNPs resulted in (rs30302002) I461V, (rs30323949) V620I and (rs30323946) S643N amino acid substitutions. However, using PolyPhen2 all three substitutions were predicted to be benign and not impact *Cog6* function [37].

### Lhfp is regulated by local eQTL in liver and bone

We next determined if the same SNPs associated with BMD regulated the expression of *Lhfp* and *Cog6*. We first searched for local expression quantitiative trait loci (eQTL) using expression data for *Lhfp* and *Cog6* in liver, brain, adipose and muscle tissues in the BXH F2 intercross. We observed a highly significant local eQTL for *Lhfp* (LOD=19.9) in liver (Figure 4A). *Cog6* was not regulated by a local eQTL in any tissue (data not shown). The lead *Lhfp* eQTL SNP (rs3665396) was located in the first intron of *Lhfp* and B6 alleles of rs3665396 were associated with increased expression of *Lhfp* relative to C3H alleles (Figure 4B). In liver we observed a negative correlation (r=−0.29, P=1.5 x 10^−4^) between *Lhfp* and BMD, as would be expected given that B6 alleles of rs3665396 were associated with decreased BMD and increased *Lhfp* (Figure 4C).

**Figure 4.**
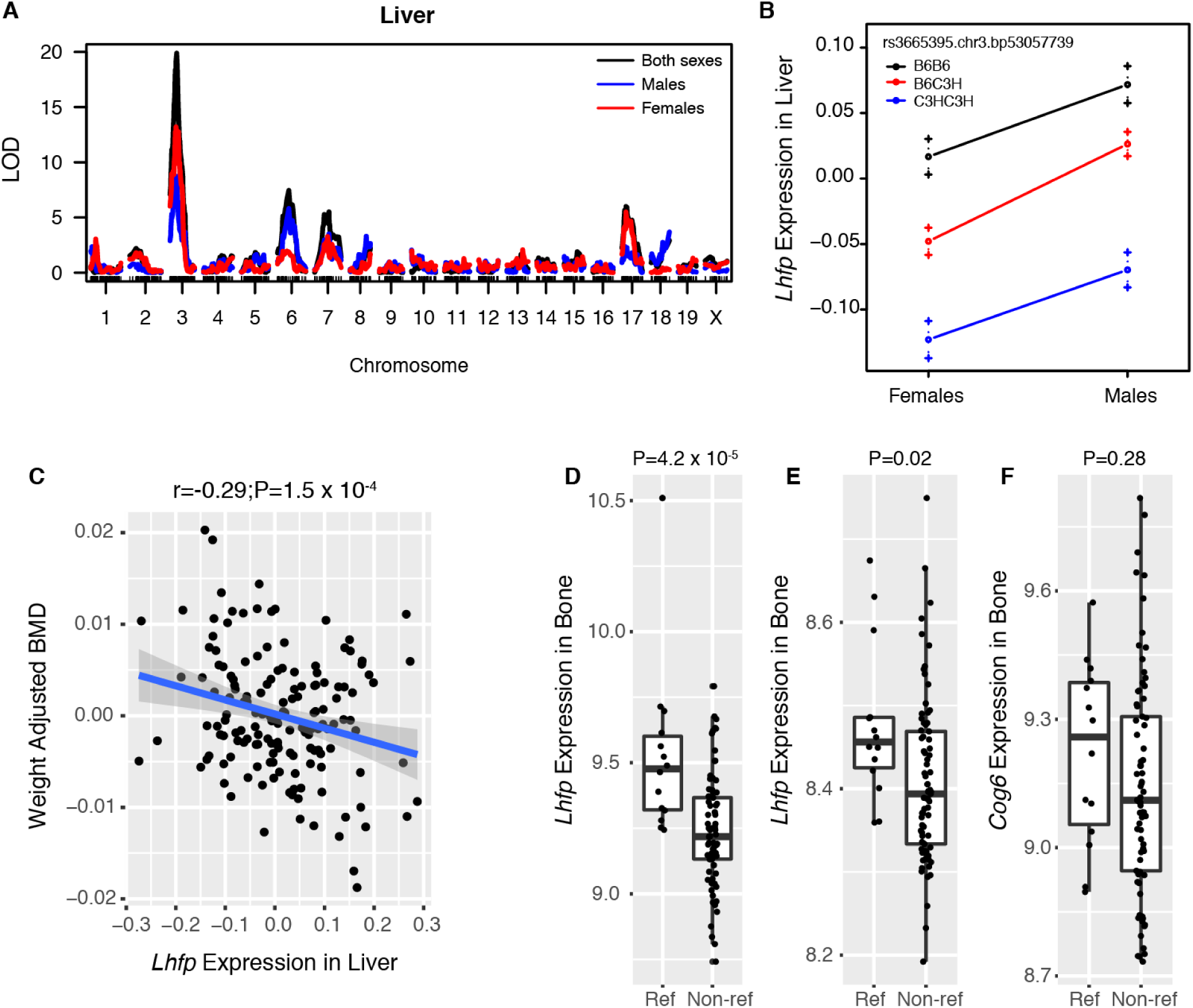
***Lhfp*** **expression is regulated by local eQTL in liver and bone overlapping the Chr.3@52.5 Mbp BMD association.** A) Strong local eQTL for Lhfp in liver tissue from BXH F2 mice. B) In BXH livers, C57BL6/J (B6) alleles at the Chr. 3 local eQTL are associated with increased *Lhfp* expression. Data are presented as means ± s.e.m. C) Lhfp expression in liver is negatively correlated with weight-adjusted femoral BMD in BXH F2 mice. D-E) *Lhfp* expression in bone in the Hybrid Mouse Diversity Panel (HMDP), measured by two independent probes (D & E), is associated with genotype of the SNP most significantly associated with BMD in the “Ackert” panel. As in the BXH F2, B6 alleles are associated with increased *Lhfp* expression in bone. F) *Cog6* expression is not associated with genotype of the SNP most significantly associated with BMD in the “Ackert” panel.

We next evaluated the expression of *Lhfp* and *Cog6* in whole bone in 95 strains of the hybrid mouse diversity panel (HMDP) [12,38]. Similar to the local eQTL in BXH F2 mice, when strains were grouped based on genotype at rs3691451 (one of the 17 peak SNPs) we observed a significant (two probes; P=4.2 x 10^−5^ and P=0.02) increase in *Lhfp* expression in strains homozygous for reference alleles (Figure 4D-E). There was no change in *Cog6* expression (P=0.28) (Figure 4F). These data are consistent with *cis* regulation of *Lhfp* in multiple tissues, including bone, suggesting that the Chr3@52.5 Mbp BMD association may act by altering *Lhfp* expression.

### Network analysis predicts a role for Lhfp in the regulation of osteoblast activity

Our group and others [12,28,32,39] have shown that co-expression network analysis can identify interactions among genes and knowledge of these interactions can assist in predicting gene function and/or the cell type in which a gene is operative. Therefore, we next used a bone co-expression network analysis to further evaluate *Lhfp* and *Cog6*. For the analysis we used a previously generated whole bone (femur with marrow removed) co-expression network from the HMDP that consisted of 13,759 genes partitioned into 21 co-expression modules [38]. In this network, *Lhfp* was a member of module 9 and *Cog6* was a member of module 2. Module 2 was enriched in a large number of gene ontology terms including “mitochondrion”, “oxidative phosphorylation” and “actin cytoskeleton”; all of which are important to bone. However, module 2 did not have a signature of a particular bone cell-type, nor was it enriched for genes known to influence BMD. In contrast, we have previously demonstrated that module 9 is enriched for genes 1) directly involved in osteoblast differentiation, 2) implicated by BMD GWAS, and 3) when knocked-out in mice impact BMD [32,40].

To investigate specific network connections for *Lhfp* and *Cog6*, we identified the 150 genes most strongly connected to each gene in their respective module. The genes with the strongest connections to *Cog6* were enriched for genes involved in “muscle structure development” (FDR=2.9 x 10^−10^), “muscle cell development” (FDR=4.4 x 10^−10^), among many other similar muscle-related categories (**Supplemental File 3**). In contrast the genes with the strongest connections to *Lhfp* were, similar to module 9, enriched for genes involved in “ossification” (FDR=1.5 x 10^−7^), “osteoblast differentiation” (FDR=8.0 x 10^−4^), “skeletal system development” (FDR=6.4 x 10^−3^), “bone development” (FDR=3.3 x 10^−2^), among many other related bone-related functional categories (**Supplemental File 4 and Figure 5A**). The *Lhfp*-centric network contained a number of genes with key roles in osteoblast differentiation and activity, including *Sp7, Pthr1, Akp2, Tmem119*, and *Bmp3* (Figure 5B). Together, these data suggest that *Lhfp* is involved in the activity of osteoblasts, a process of direct relevance to the regulation of bone mass.

**Figure 5.**
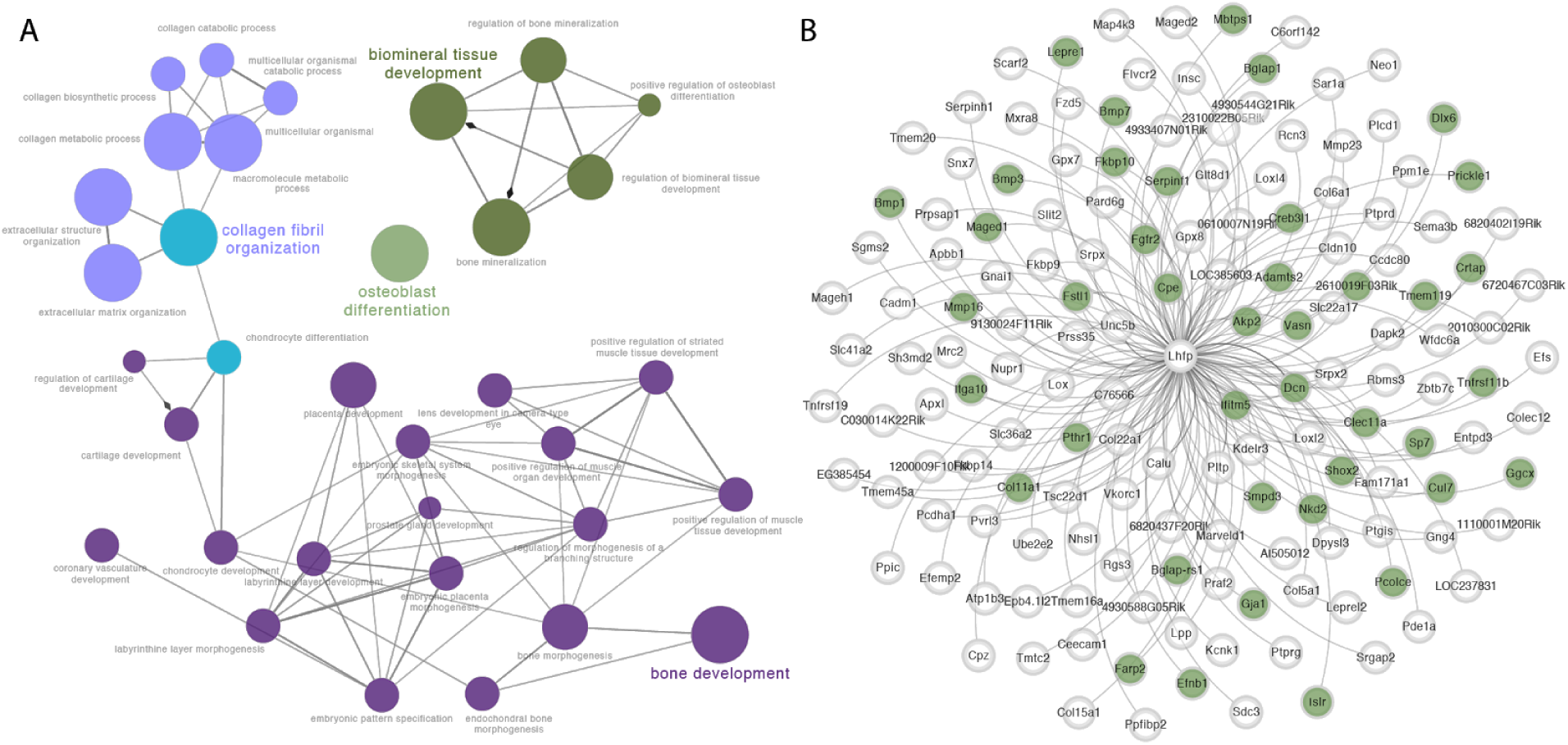
***Lhfp*** **in bone is highly connected to genes involved in osteoblast differentiation**. A) Network depiction of gene ontology “biological process” categories containing more of the genes with strong connections to Lhfp in a bone co-expression network than would be expected by chance. B) Genes with the strongest connections to Lhfp in a bone co-expression network. The genes highlighted in green have been shown to be directly involved in osteoblast differentiation.

### Lhfp regulates the number and osteogenic differentiation of bone marrow stromal cells

Bone marrow stromal cells (BMSCs) are adherent marrow cells that contain the mesenchymal progenitors of osteoblasts [41]. To test the role of *Lhfp* in osteoblast function, we quantified the number of BMSCs and their ability to form osteoblasts from mice lacking *Lhfp*. Using CRISPR/Cas9, we created five small deletions (ranging from 4-16 bps) in exon 2 (ATG start codon is in exon 2) of *Lhfp* (Table 1). All five are nonsense mutations resulting in a truncated LHFP protein. As expected, we observed significantly decreased *Lhfp* transcript levels in heterozygotes (*Lhfp*^+/−^) and mutants (*Lhfp*^−/−^) from all five lines (Figure 6A). Since all five mutations impacted *Lhfp* expression in the same manner, we grouped littermate mice by genotype from all lines for all downstream experiments.

**Table 1.**
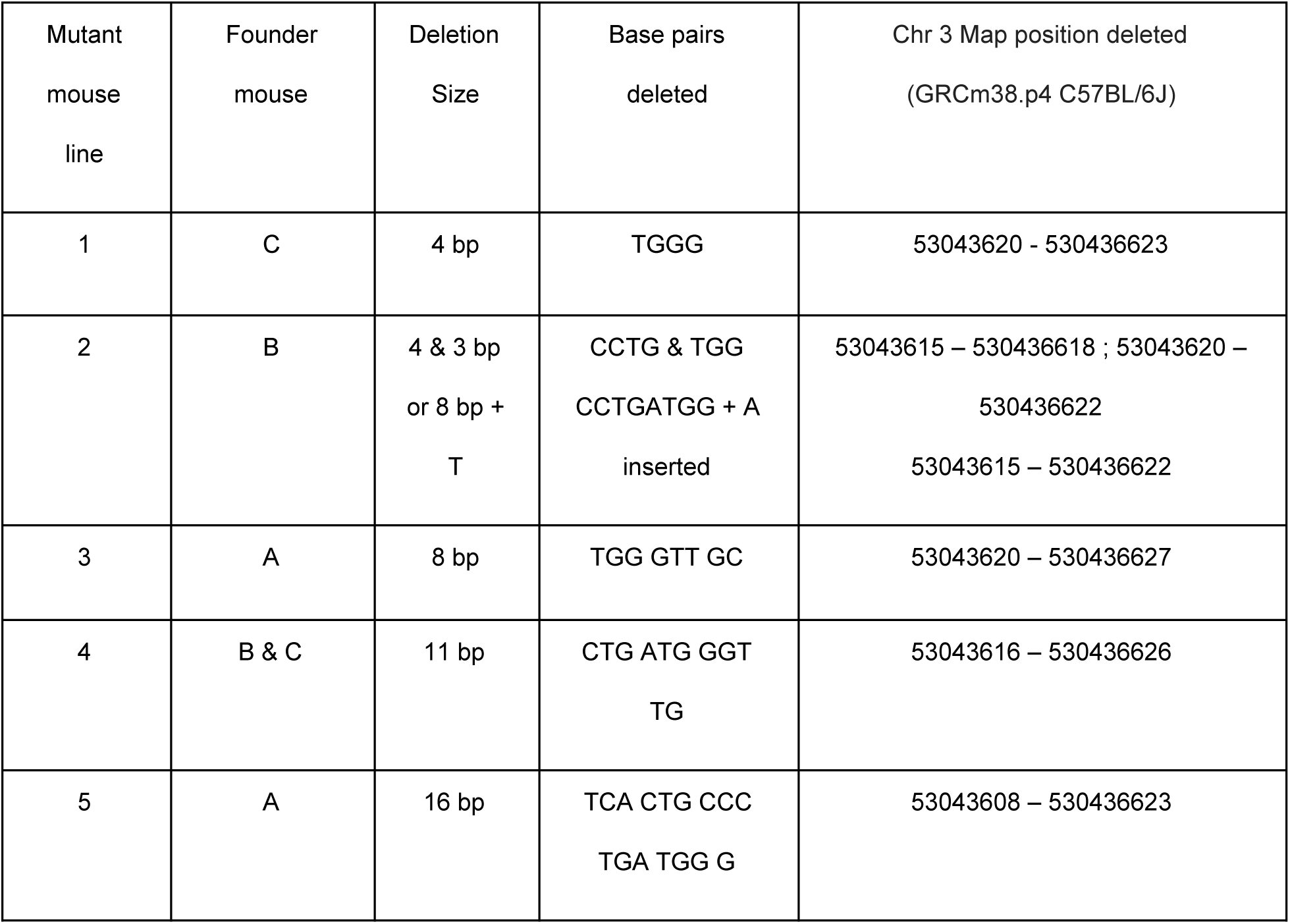
**Description of CRISPR/Cas9-induced** ***Lhfp*** **mutations.**

**Figure 6.**
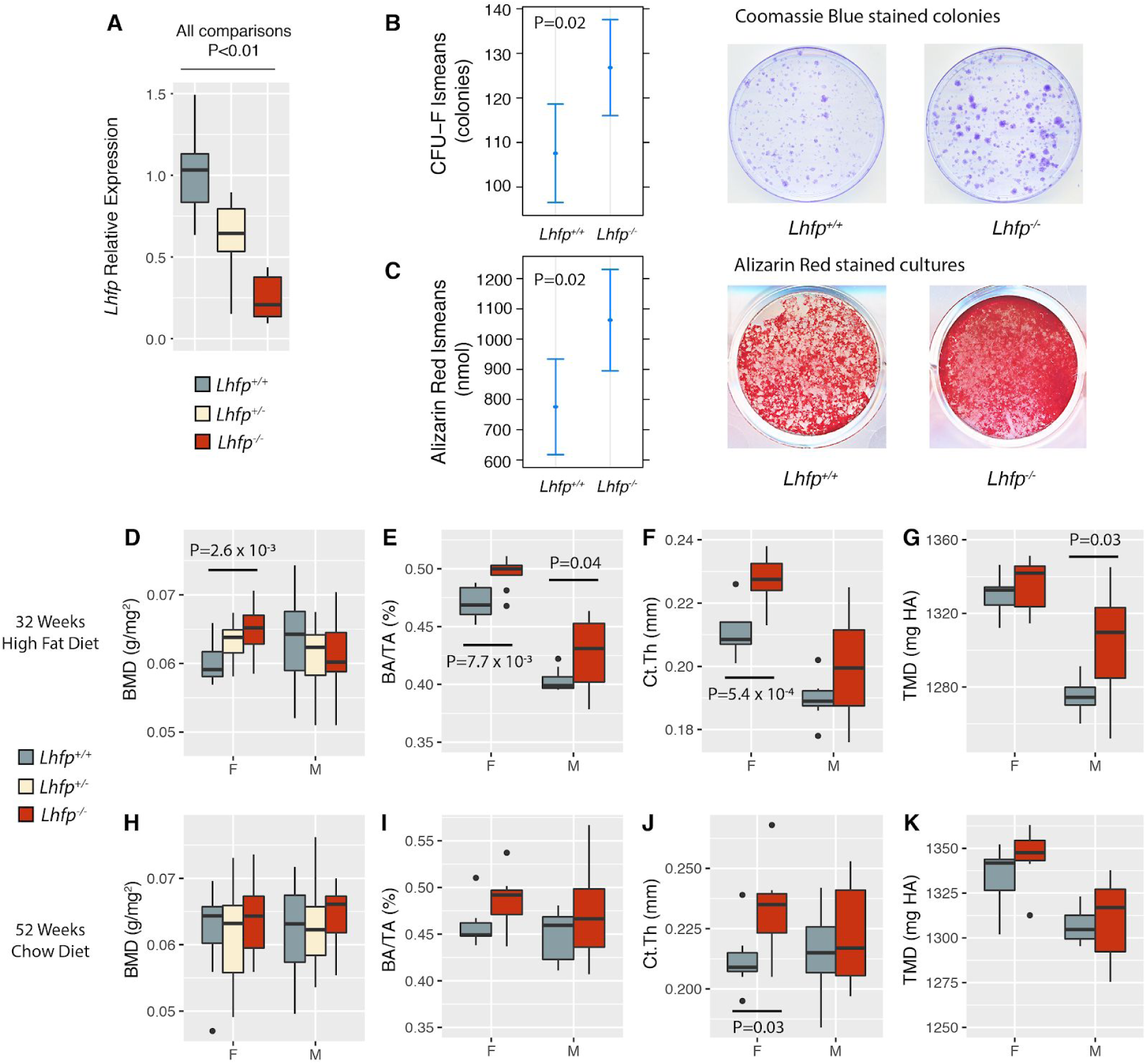
***Lhfp*** **is a negative regulator of osteoblast differentiation and cortical bone mass.** A) *Lhfp* transcript levels are decreased due to the five nonsense mutations created using CRISPR/Cas9 (*Lhfp*^+/+^, N=10; *Lhfp*^+/−^, N=11; *Lhfp*^−/−^, N=7; represents at least one mouse from each of the five mutant lines, see Table 1). Data are presented as means ± 1.5 times interquartile range (IQR). B) CFU-F colonies are increased in Lhfp-/- mice (*Lhfp*^+/+^, N=41; *Lhfp*^−/−^ N=43). Data are presented as means ± s.e.m. Images are from the sample closest to the mean for each genotype. D-G) Femoral BMD, cortical bone area fraction (BA/TA), cortical thickness (Ct.Th), and tissue mineral density (TMD) in Lhfp mutant mice in 32 week old mice fed a high-fat diet. Data are presented as means ± 1.5 times IQR. H-K) Femoral BMD, cortical bone area fraction (BA/TA), cortical thickness (Ct.Th), and tissue mineral density (TMD) in Lhfp mutant mice in 52 week old mice fed a chow diet. Data are presented as means ± 1.5 times IQR.

Next, we performed colony-forming unit-fibroblast (CFU-F) assays, a direct measure of BMSCs, in 16 week-old *Lhfp*^−/−^ and littermate *Lhfp*^+/+^ mice. We observed similar trends in both sexes; therefore, all data were combined and adjusted for the effects of sex to increase power. In *Lhfp*^−/−^ mice, we observed a significant (P=0.02) increase in CFU-F number (Figure 6B). We next evaluated the ability of BMSCs from *Lhfp*^+/+^ and *Lhfp*^−/−^ mice to differentiate into mineralizing osteoblasts. Consistent with network predictions, *Lhfp*^−/−^ BMSCs exhibited increased mineralization as measured by bound alizarin red (P=0.02; Figure 6C) and percent of mineralized surface (P=0.009; data not shown).

### Lhfp regulates cortical bone mass

We next determined if bone mass was altered in *Lhfp* mutant lines. To replicate the conditions of the Ackert inbred strain panel and the BXH F2, we generated two cohorts of mice. The first was fed a chow diet for 12 months, while the second was fed a high-fat diet from 8 to 32 weeks of age. Based on the negative correlation between BMD and *Lhfp* expression, the direction of the genetic effects on expression in bone, and increased osteoblast activity observed above, we predicted increased BMD in *Lhfp*^−/−^ mice. In both cohorts, BMD was measured in mice of all three genotypes and cortical and trabecular microarchitecture was measured by microCT only in *Lhfp*^+/+^ and *Lhfp*^−/−^ mice. At 32 weeks of age in mice on a high-fat diet we observed significantly (P=2.6 x 10^−3^) increased femoral BMD as a function of mutant *Lhfp* alleles in females, but not males (Figure 6D).

BMD is an inherently noise phenotype; therefore, to generate a more detailed understanding of the effects of *Lhfp* in bone we used microCT to investigate the amount of bone in both the femoral trabecular and cortical compartments. We did not observe effects on trabecular bone mass at the distal femur in either male or female mice (data not shown). However, *Lhfp*^−/−^ mice of both sexes had significantly (P<0.05) increased femoral cortical bone area fraction (BA/TA) and cortical thickness (Ct.Th) as compared to *Lhfp*^+/+^ littermates (Figure 6E-F). We also observed a significant (P=0.03) increase in tissue mineral density (TMD) in male *Lhfp*^−/−^ mice (Figure 6G). In general, we observed the same trends of increased cortical bone mass in *Lhfp*^−/−^ mice at 52 weeks of age; however, only Ct.Th in females was significant (P=0.03) (Figure 6H-K). These data indicate that *Lhfp* is a negative regulator of cortical bone mass in both male and female mice. They are also consistent with *Lhfp* underlying, at least in part, the BMD association on Chr. 3@52.5 Mbp.

## DISCUSSION

In this study, we used GWAS in a mouse inbred strain panel and a multifaceted systems genetics approach to identify and validate a high-resolution association for BMD on Chr. 3. The association directly implicated four genes: *Gm2447, Gm20750, Cog6* and *Lhfp*. Of these, *Lhfp* expression was regulated by a local eQTL in liver and bone and was predicted, based on a bone gene co-expression network, to be involved in osteoblast-mediated bone formation. We demonstrated that mice deficient in *Lhfp* displayed increased BMSC number and increased BMSC osteogenic differentiation. Furthermore, *Lhfp*^−/−^ had increased BMD due to increased cortical bone mass. Together these data indicate that *Lhfp* is responsible, at least in part, for the BMD association we identified on Chr. 3 at 52.5 Mbp. This work defines *Lhfp* as a negative regulator of the pool of osteoprogenitor cells, osteoblast activity, and cortical bone mass.

GWAS in mice has proven to be a powerful approach for the identification of genomic regions harboring trait-associated genetic variation [11]. The earliest applications of GWAS in mice used panels of readily accessible inbred strains [18–20]. However, such approaches were plagued by false positives due to population stratification [23]. Aware of this limitation, we first performed GWAS for BMD after correcting for population structure in inbred strains and then replicated the analysis in two seperate strain panels (containing many of the same strains, but representing independent measures of BMD in different environments) and an F2 intercross. Of the multiple loci identified, the association on Chr. 3 at 52.5 Mbp was identified in all datasets, strongly suggesting it represents a bona-fide genetic association.

The Chr. 3 locus, as defined by the interval harboring the most significant SNPs, contained four genes; *Gm2447, Gm20750, Cog6* and *Lhfp. Gm2447* and *Gm20750* were both predicted lncRNAs. This prediction is based on limited data and the fact that we did not observe their expression in bone tissue or osteoblasts, suggest they are not likely causal for the locus; though, this alone is not enough to definitely exclude their involvement. For *Cog6* and *Lhfp* we used eQTL data and a bone co-expression network to assist in evaluating their potential causality. Both analyses supported a role for *Lhfp*. Using eQTL data from liver tissue in the BXH F2 intercross and bone tissue from the HMDP, we observed that variants associated with decreased BMD were associated with increased expression of *Lhfp*. We did not observe an association between the BMD-associated variants and *Cog6* expression. Furthermore, *Lhfp* was a member of a well-studied module of co-expressed genes in mouse bone. This module is highly enriched for genes that play a role in osteoblast function, which provides a direct explanation as to how *Lhfp* may be impacting BMD. In contrast, *Cog6* was a member of a module enriched for genes involved in a wide range of “energy-generating” functions. Importantly, all of our experimental results confirmed that *Lhfp* is a negative regulator of osteoblast activity and BMD. While these data support a role for *Lhfp* in the effects of the Chr. 3 locus, they do not exclude any of the other genes in, as well as flanking the locus.

In all four genetic populations used to identify the association on Chr. 3@52.5 Mbp, the strength of the association differed by sex. For example, in the “Ackert” population the association was stronger in males relative to females. In the “Naggert” strain set the strength of the association was similar in both sexes, albeit both were lower than seen in the other three populations. Similar to the “Ackert” strains, the association was stronger in males than females in the “Tordoff” strain set. In the BXH F2, the Chr. 3 QTL was male-specific, with little to no signal in females. The increase in cortical bone mass in *Lhfp*^−/−^ mice was also sexually dimorphic. Although *Lhfp* deficiency increased cortical bone mass in both sexes in general, the effects were slightly more pronounced in females as compared to males. The discrepancies could be the result of inaccuracies in estimating genetic effect sizes in the relatively small strain sets, or influences from the different genetic backgrounds of the populations studied (strains sets vs. F2 vs. knockout).

Little is known regarding the molecular function of *Lhfp*. *Lhfp* is a member of the *Lhfp-like* gene family, which is a subset of the superfamily of tetraspan transmembrane protein encoding genes. It was first identified as a translocation partner with the HMGIC gene in benign lipomas [35]. Since its initial discovery it has been implicated in the mesenchymal differentiation of gliosarcoma [42]. The human *LHFP/COG6* locus was also identified by GWAS as harboring variants associated with hippocampal volume [43]. However, prior to this study *Lhfp* had not been connected to the regulation of osteoblast function or BMD. Based on our experimental results, we hypothesize that *Lhfp* regulates bone mass by increasing the number and osteogenic potential of mesenchymal stem cells. However, further work will be required to elucidate its precise molecular role in osteoblasts and bone.

In summary, we have used GWAS in a set of inbred strains to identify an association impacting femoral BMD on Chr. 3 at 52.5 Mbp. We show using a variety of approaches that *Lhfp* is a causal gene and likely responsible for most, if not all, of the effects of this locus. Our results identify *Lhfp* as a novel negative regulator of osteoblast function and BMD and increase our understanding of the genetics of BMD.

## ACKNOWLEDGEMENTS

The authors would like to acknowledge grant support from the National Institute of Arthritis and Musculoskeletal and Skin Disease (R01AR057759 to CRF and R21AR060981 to CLA) and the National Institute of General Medical Sciences (R01GM070683 for GC and DG) of the National Institutes of Health. We also thank Dr. Wenhao Xu in the Genetically Engineered Murine Model (GEMM) Core at the University of Virginia (UVa) for assistance in generating the *Lhfp* CRISPR/Cas9 mutants. Partial funding for generation of the *Lhfp* mutant mice was provided by the UVa Cancer Center (P30CA044579) and microCT analysis was partially fund through the Biomechanics, Biomaterials, and Multimodal Tissue Imaging Core in the UR Center for Musculoskeletal Research (P30AR069655).

## MATERIALS AND METHODS

### Association Analysis

The “Ackert” strain set contained BMD data on 32 inbred strains at three time points (6, 12 and 18 months). In the present study, we focused on BMD measures at 12 months of age. The complete dataset is available from the Mouse Phenome Database (MPD) (https://phenome.jax.org/projects/Ackert1). After removing wild-derived strains, and C57BLKS/J (due to inclusion of this strain producing spurious results) we were left with data on 27 strains. For the association analysis BMD was rankZ transformed. SNPs were obtained from strains genotyped on the Mouse Diversity Array (http://churchill-lab.jax.org/website/MDA) [44]. SNPs with a minor allele frequency < 0.05 were removed, leaving 228,085 SNPs. These SNPs were used to generate a kinship using the ‘emma.kinship’ R script available in the Efficient Mixed Model Association (EMMA) R package (available at http://mouse.cs.ucla.edu/emma/) [24]. The emma.REML.t function of EMMA was used to perform all mapping analyses. The significance of the maximum association peak was assessed by performing 1,000 permutations of the data. In each permutation, the minimum p-value was recorded to produce an empirical distribution of minimum permutation p-values. The quantiles of this distribution were used to assign adjusted p-values. GWAS resulted were visualized using the “qqman” R package [45]. Replication of the association on Chr. 3@52.5 Mbp was performed using “Tordoff” and “Naggert” inbred strains sets. These data are available from MPD (https://phenome.jax.org/projects/Naggert1 and https://phenome.jax.org/projects/Tordoff3). Replication analyses were restricted to Chr. 3 and otherwise performed as described above.

### Generating expression profiles of *Cog6* and *Lhfp*

Femora were isolated from an inbred Collaborative Cross strain (CC016/GeniUnc; Jackson Lab Stock #024684) (N=3 mice). Marrow was isolated and bone marrow stromal cells (BMSCs) were differentiated as described below. Total RNA was then isolated from bone and BMSC-derived osteoblasts using RNeasy Plus Mini Kit (Qiagen) [28]. RNA-Seq libraries were constructed using TruSeq RNA Library Prep Kit v2 sample prep kits (Illumina). Samples were sequenced to an average depth of 24.6 million 2 x 75 bp paired-end reads on an Illumina NextSeq500 sequencer. Fastq files were aligned to the mouse reference (GRCm38) using HISAT2 v 2.0.5 (https://ccb.jhu.edu/software/hisat2/index.shtml) with a SNP aware reference index (genome_snp) [46]. Expression levels in Fragments Per Kilobase of transcript per Million mapped reads (FPKM) were generated using Stringtie [47]. The data are available from GEO (GSE121887). Microarray profiles for *Cog6* and *Lhfp* in 96 tissues/cell-types were downloaded from BioGPS (http://biogps.org).

### EQTL analyses

The generation of microarray expression data and eQTL analyses on bone from the 96 strains of the Hybrid Mouse Diversity Panel (HMDP) has been previously described [7]. These data are available from NCBI Gene Expression Omnibus (GEO) (GSE27483). A t-test was used to test for differences in *Cog6* and *Lhfp* expression in strains stratified by genotype at rs3691451 (of the 17 peak BMD SNPs). Liver, brain, muscle and adipose eQTL in the BXH F2 were identified using R/qtl [48]. The expression data is available from GEO (GSE11338, GSE11065, GSE12798, and GSE12795). The genotypes and expression data are also available from GeneNetwork (“BH/HB F2 UCLA”, http://www.genenetwork.org/webqtl/main.py).

### Network Analysis

The generation of a bone co-expression network and characterization of the module 9 (M9) is described in [40,48]. We identified genes with the strongest connections to *Cog6* and *Lhfp* based on Topological Overlap Measures (TOMs), calculated as described in [31]. Network depictions were constructed using Cytoscape [24]. Gene Ontology (GO) analysis was performed using the PANTHER database statistical overrepresentation test (http://www.pantherdb.org/) [49]. The analysis was restricted to the “GO biological process complete” annotation data set.

### Generation of Lhfp mutant mice

The *Lhfp* knockout mice used in this study were generated using the CRISPR/Cas9 genome editing technique. Cas9 mRNA that was injected into C57BL6/N embryos was synthesized exactly as outlined in [50] while the guide RNA (sgRNA) was generated with some modifications. Briefly, the 20 nucleotides (nt) sequence that would be used to generate the sgRNA was chosen using the CRISPR design tool developed by the Zhang lab (crispr.mit.edu). The chosen sequence and its genome map position is homologous to a region in Exon 2 that is approximately 300 bp, 3’ of the start codon (the ‘ATG’ is located in Exon 2 of *Lhfp*) (Supplemental Table 1). To generate the sgRNA that would be used for injections oligonucleotides of the chosen sequence, as well as the reverse complement (Supplemental Table 1, primer 1 and 2, respectively), were synthesized such that an additional 4 nts (CTTC and AAAC) were added at the 5’ ends of the oligonucleotides for cloning purposes. These oligonucleotides were annealed to each other by combining equal molar amounts, heating to 90°C for 5 min. and allowing the mixture to passively cool to room temperature. The annealed oligonucleotides were combined with BbsI digested pX330 plasmid vector (provided by the Zhang lab through Addgene; https://www.addgene.org/) and T4 DNA ligase (NEB) and subsequently used to transform Stbl3 competent bacteria (Thermo Fisher) following the manufacturer’s’ protocols. Plasmid DNAs from selected clones were sequenced from primer 4 (Supplemental Table 1) and DNA that demonstrated accurate sequence and position of the guide were used for all downstream applications. The DNA template used in the synthesis of the sgRNA was the product of a PCR using the verified plasmid DNA and primers 3 and 5 (Supplemental Table 1). The sgRNA was synthesized via in vitro transcription (IVT) by way of the MAXIscript T7 kit (Thermo Fisher) following the manufacturer’s protocol. sgRNAs were purified and concentrated using the RNeasy Plus Micro kit (Qiagen) following the manufacturer’s protocol.

C57BL/6N female mice (Envigo) were super-ovulated and mated with C57BL/6N males. The females were sacrificed and the fertilized eggs were isolated from the oviducts. The fertilized eggs were co-injected with the purified Cas9 mRNA (100 ng/μl) and sgRNA (30 ng/μl) under a Leica inverted microscope equipped with Leitz micromanipulators (Leica Microsystems). Injected eggs were incubated overnight in KSOM-AA medium (Millipore Sigma). Two-cell stage embryos were implanted on the following day into the oviducts of pseudo pregnant ICR female mice (Taconic or Envigo). Pups were screened by PCR of tail DNA using primers 6 and 7 with subsequent sequencing of the resultant product from primer 8 (Supplemental Table 1).

Two sets of injections (of ∼100 eggs each) were performed resulting in 2 mice possessing mutations from each set of injections (A,B and C,D, respectively; Table 1). All 4 mice possessed out of frame bi-allelic deletions ranging from 1-16 bp; progeny from only 3 of the founders (mice A, B, C, Table 1)) were used in this study. Note that an identical 11bp deletion was found in two mice from two separate injections. qPCR with primers 9 and 10 (Supplemental Table 1) was used to assess *Lhfp* expression as outlined in [32].

### CFU-F and osteogenic differentiation assays

Isolation [51] and differentiation of mesenchymal stromal cells (MSC) from the bone marrow of mouse femurs was performed as described for osteoblasts [52] with minor modifications. Briefly, one or both femurs from a given mouse were aseptically isolated, denuded of soft tissue and the marrow extracted by removing the proximal end of each bone and centrifuging at 2000 xg for 30 s such that the marrow collects into 25 µl of fetal bovine serum (FBS). Exudates from a single femur were dispersed, via trituration, in 5ml complete media (MEM-alpha, 10% FBS, 100U penicillin/100ug Streptomycin per ml, 2 mM glutamine). Cells were manually counted where upon 4 million were used to seed a 10 cm dish for CFU-F determination and the remainder were applied to a 60mm dish. Media on the 10 cm dishes was changed on days 2, 4 and 8; on day 14, cells were fixed (NBF), stained (Coomassie, BioRad #161-0436) and the number of CFU-Fs determined via Image J analysis and the manual counting of colonies. Media on the 60 mm dishes was changed on day 2 with cells removed via trypsin/EDTA (Gibco) digestion on day 4. Detached cells were triturated in 5ml complete media, pelleted at 1000 xg for 5 min., re-suspended in 1 ml complete media and counted where upon 150,000 cells were used to seed a well of a 12 well dish. A minimum of 2 wells per sample were obtained for all samples reported here in. Osteoblast differentiation was initiated 3 days after plating (7 days after bone marrow isolation) by replacing the media with complete media supplemented with 50 µg/ml ascorbic acid, 10 mM beta-glycerophosphate and 10 nM dexamethasone. Media was changed every other day for 8 days at which time cells were either used as a source for RNA (mirVana, Thermo Fisher) or used to determine the amount of hydroxyapatite formed during differentiation [32]. Briefly, cells were washed with PBS, fixed with neutral buffer formalin (NBF) for 15 min. and subsequently stained with 40 mM Alizarin Red (AR), pH 5.6 for 20 min and washed extensively with H_2_O. The amount of AR bound to mineral was quantitated by Image J analysis of scanned images as well as the 5% Perchloric Acid eluate absorbance at 405 nm.

### Measurement of BMD and microarchitecture (microCT)

Femoral BMD was measured using a Lunar PIXImus II Mouse Densitometer (GE Medical Systems Model 51045; Madison, WI, USA). Morphologies of the trabecular bone of the distal femur and cortical bone of the femoral midshaft were measured using micro-focus X-ray computed tomography (vivaCT 40, Scanco Medical AG, Bassersdorf, Switzerland) following guidelines for assessment of bone microstructure [53]. Tomographic volumes were acquired at 55 kV and 145 µA, collecting 2000 projections per rotation at 300 millisecond integration time. Three-dimensional 16-bit grayscale images were reconstructed using Scanco Evaluation software, Version 6.5-3. Threshold values were adjusted to best match the silhouette of features of interest in the threshold-subtracted image compared to the grey-scale image. The resulting threshold for hydroxyapatite-equivalent density was 370 mg/cm^3^ for compact or cortical bone and 270 mg/cm^3^ for the trabecular bone region; these values were applied to subsequent samples. Volumetric analysis was confined to the trabecular region for the distal femur by manual exclusion of the cortical bone. A 1.03 mm high region of interest was analyzed beginning at 1 mm proximal to the growth plate. For the cortical bone, a 0.3 mm high region was analyzed at the mid-diaphysis. Measures analyzing the distal femur trabecular site included total volume, bone volume, trabecular bone volume fraction (BV/TV), thickness, number, connectivity density and structure model index (SMI). Cross-sectional measurements of the cortical bone included bone volume, total volume, marrow area and polar moment of inertia.

### Statistical analysis of Lhfp mutant data

All statistical analyses were conducted using the R language and environment for statistical computing [54]. The *Lhfp* qPCR data was analyzed using a t-test. Data are presented as means ± 1.5 times the interquartile range.

CFU-F and osteogenic differentiation data was analyzed using the “lsmeans” R package [55]. The data were fit to a linear model including the effects of genotype and sex. P-values were adjusted using the “tukey” method [55]. Data are presented as lsmeans ± s.e.m. BMD and microarchitectural bone data were analyzed using ANOVA with a linear model including the effects of genotype, body weight, and any other phenotype-specific covariates. Data are presented as means ± 1.5 times the interquartile range.

